# A New Sparse Bayesian Quantile Neural Network-based Approach and Its Application to Discover Physiological Sweet Spots in the Canadian Longitudinal Study on Aging

**DOI:** 10.64898/2026.02.19.706407

**Authors:** Joosung Min, Olga Vishnyakova, Angela Brooks-Wilson, Lloyd T. Elliott

## Abstract

Identifying physiological sweet spots (optimal ranges for homeostasis) is essential for precision medicine. However, traditional statistical methods often rely on globally linear or locally jagged models that struggle to capture the smooth, non-linear nature of biological regulation in high-dimensional data. We present the *Quantile Feature Selection Network* (Q-FSNet), a neural network-based framework that integrates quantile regression, feature selection, and uncertainty estimation to identify biomarkers with sweet spots. Unlike traditional methods, Q-FSNet learns continuous response curves without requiring pre-specified number of change points. We further introduce *Quantile Dirichlet Network* (Q-DirichNet), a fully Bayesian extension that utilizes Dirichlet priors to automate feature shrinkage. Using data from the Canadian Longitudinal Study on Aging, we identified 25 metabolites with distinct homeostatic ranges for which biological age acceleration is minimized. The metabolites with sweet spots for biological aging include some derived from diet or produced by the gut microbiome; this highlights their potential for knowledge translation and public health impact. Our results, corroborated by existing literature, demonstrate that these sparse neural network-based methods offer a scalable and interpretable tool for discovering metabolic signatures of healthy aging vs. dysregulation in large-scale omics research.

## 1 Introduction

Precision medicine aims to tailor prevention and treatment strategies to an individual’s biological profile. A central idea underlying this goal is *homeostasis*: the regulation of physiological systems to maintain internal stability. Many biological markers operate optimally within specific ranges, or physiological “sweet spots” [1]. When these ranges are maintained, biological systems function well. Well-known examples include vitamin D levels linked to calcium homeostasis and mortality [2,3], as well as fasting glucose levels associated with cardiovascular risk [4]. More broadly, deviations from physiological sweet spots reflect loss of regulatory control and are implicated in metabolic disorders that become increasingly common with aging [1,5,6].

Identifying such sweet spots in large-scale molecular data remains challenging. Previous work has relied on segmented (or piecewise) regression to detect threshold effects [1,2,7]. While useful, these approaches require the number and location of change points to be specified in advance and typically assume abrupt changes in slope, yielding a non-differentiable objective function. These assumptions may not reflect biological reality, where regulation is often smooth and adaptive. In addition, fitting separate models for each biomarker limits scalability and does not leverage shared information across metabolites.

These limitations are particularly relevant for age-related phenotypes, for which risk is often concentrated at the extremes of the outcome distribution rather than near the mean. Age acceleration, defined as the difference between biomarker-derived age estimates (such as HannumAge [8], PhenoAge [9] and GrimAge [10]) and chronological age, captures this heterogeneity and is strongly associated with morbidity and mortality. Individuals with accelerated aging may exhibit metabolomic profiles that differ substantially from those aging more slowly. Understanding these subgroup-specific metabolic signatures is essential for identifying dysregulation and potential intervention targets.

Conventional regression methods focus on average effects and therefore obscure how metabolite– phenotype associations vary across the outcome distribution. Quantile regression [11] addresses this limitation by modeling effects at specific quantiles, allowing relationships to differ between low-, middle-, and high-risk groups. This makes quantile regression particularly well suited for studying biological heterogeneity. However, quantile regression has seen limited use in high-dimensional settings such as metabolomics, wherein non-linear relationships, correlated features, and heteroscedastic noise are common.

Traditional quantile regression methods based on machine learning approaches, including lasso [11,12] and gradient-boosted machines [13], may not be not well suited for identifying physiological sweet spots. These methods rely on piecewise linear or stepwise decision rules that inherently yield predictions that are either globally linear or locally jagged and discontinuous, making it difficult to identify optimal or dysregulated physiological ranges. Examples of quantile regression curves from different modeling approaches are illustrated in Figure 1.

**Figure 1.**
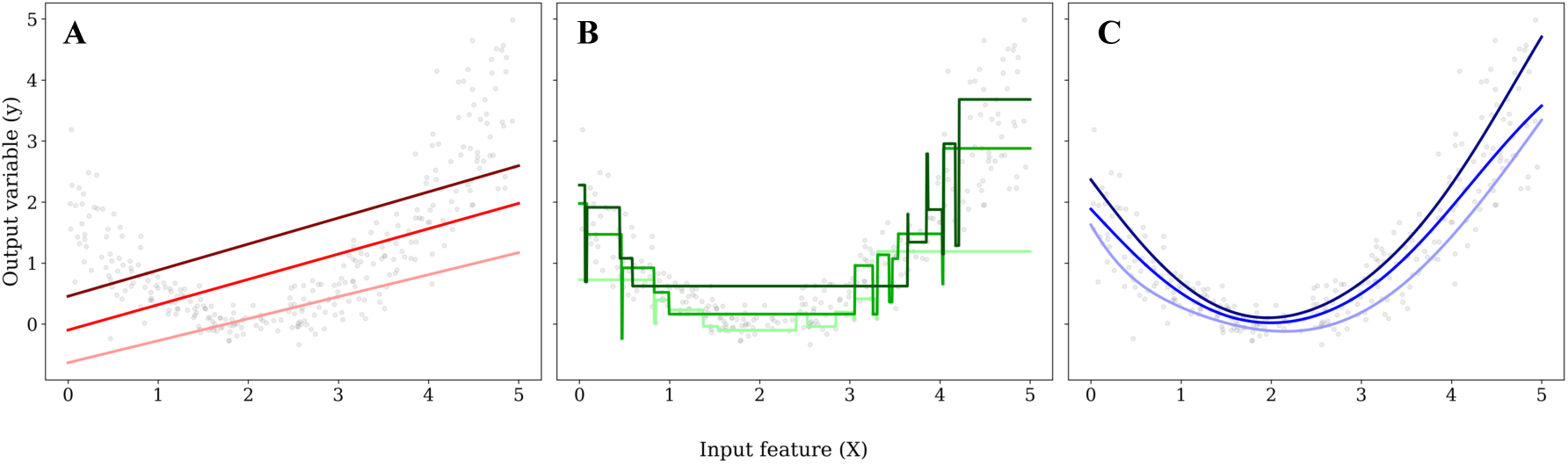
Illustration of quantile regression fits obtained using different modeling approaches. (A) Linear quantile regression, (B) Gradient-boosted machines, (C) Neural network. NN-based quantile regression produces smooth, continuous, and differentiable curves across quantiles, facilitating detection of physiological sweet spots.

Neural networks [14,15] offer the flexibility needed to model complex, non-linear associations in high-dimensional data, but they also present challenges. In high dimensional tabular settings, prevalent in biomedical datasets, standard neural networks are prone to overfitting and can be influenced by many weak or irrelevant features, especially in small sample sizes. This makes the models sensitive to noisy or uninformative predictors [16–20]. Consequently, it is often beneficial to suppress or eliminate the influence of irrelevant features through explicit feature selection or feature weighting mechanisms. These approaches can enhance both generalization and interpretability [16,19,20]. Moreover, their lack of reliable uncertainty estimates limits their use in clinical and epidemiological research, where understanding prediction confidence is critical [21–23]. Fully Bayesian neural networks address this issue but are often computationally infeasible at scale [24].

To address these challenges, we propose the Quantile-Feature Selection Network (Q-FSNet), an end-to-end framework that integrates quantile regression with neural networks, feature selection, and uncertainty estimation. By using a quantile regression loss [11], Q-FSNet directly targets specific regions of the outcome distribution, including high-risk extremes. Unlike segmented regression, this approach does not require pre-specification of the number or location of change points and does not assume abrupt threshold behavior. Instead, the model learns smooth, non-linear response curves and identifies points of inflection in a data-driven manner. A feature selection layer [25] enables joint modeling of all metabolites within a single framework, while suppressing irrelevant or weakly informative features, thereby reducing noise and improving predictive performance. In addition, we use Monte Carlo dropout layers [26] to provide practical estimates of predictive uncertainty. We also introduce a fully Bayesian extension, the Quantile Dirichlet Network (Q-DirichNet), which reformulates the feature selection process within a hierarchical Bayesian framework and provides a fully probabilistic baseline for comparison with Q-FSNet.

We applied Q-FSNet and Q-DirichNet to publicly available benchmark datasets and to high-dimensional metabolomic data from the Canadian Longitudinal Study on Aging (CLSA; [27]). While the CLSA is a longitudinal project, this study utilized the baseline comprehensive dataset, resulting in a cross-sectional design. Q-FSNet showed competitive predictive performance compared to traditional quantile regression methods by suppressing the signals from irrelevant features, while also offering predictive uncertainty. More importantly, this approach can reveal smooth, non-linear relationships between metabolites and phenotypes. By applying marginalized response functions to these patterns, we were able to estimate physiological sweet spots that might be difficult to detect using other methods. This technique isolates the marginal effect of a specific metabolite from the broader metabolic context, offering a more nuanced view of homeostatic ranges than models that assume strict linearity or produce the “staircase” artifacts common in tree-based methods.

Applying of Q-FSNet to the CLSA cohort, we examined 804 metabolites and discovered 25 metabolites with sweet spots where biological age acceleration is the lowest. These consist of 7 amino acids, 14 lipids, 3 xenobiotics, and 1 vitamin. These findings suggest that Q-FSNet can serve as a scalable tool for precision health and medicine research, balancing the inherent complexity of high-dimensional metabolomics with biological interpretability.

## 2 Results

### 2.1 Evaluation on publicly available datasets

Before applying our methods to the CLSA data, we evaluated their predictive performance on several publicly available benchmark datasets from the University of California Irvine Machine Learning Repository [28]. To mirror the sample sizes of the CLSA analyses (*N*=635 for males and *N*=653 females), we focused on datasets with relatively small numbers of observations. The benchmark datasets included Airfoil Noise (*N*=1,503, *K*=5; [29]), Boston Housing (*N*=506, *K*=13; [30]), Concrete Strength (*N*=1,030, K=8; [31]), Diabetes (*N*=442, *K*=10; [32]), Energy Efficiency (*N*=768, *K*=8; [33]), and Yacht Hydrodynamics (*N*=308, *K*=6; [34]), where K denotes the number of input features. We assessed our two proposed models against three baseline quantile regression approaches: Lasso-based quantile regression [11,12], gradient-boosted machines (GBM) [13,35], and a standard multilayer perceptron (MLP) [14,15]. Model performances were evaluated using quantile loss and empirical coverage probability, with results summarized in Figure 2.

**Figure 2.**
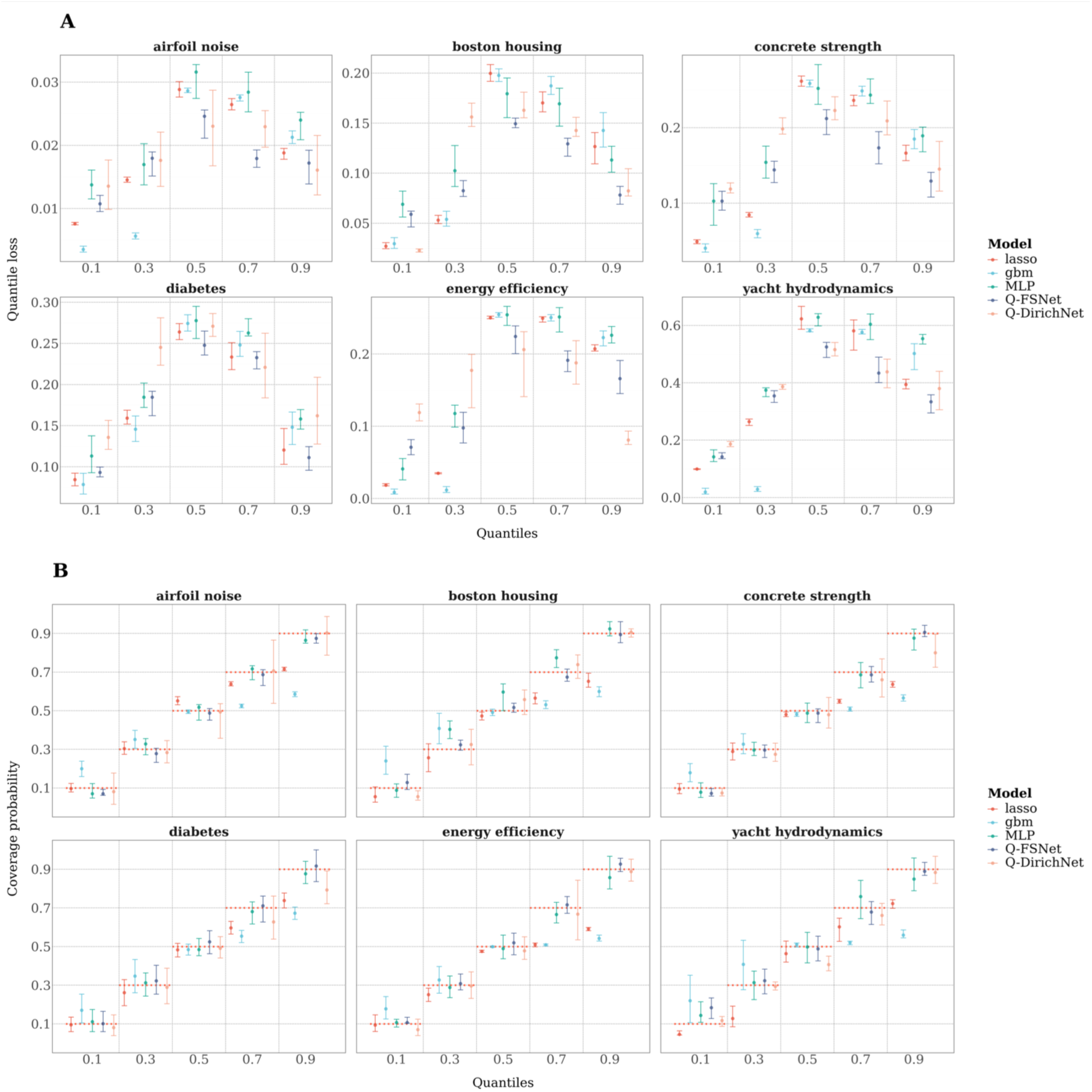
Predictive performance of quantile regression models evaluated on six publicly available benchmark datasets. lasso-based quantile regression (lasso), gradient-boosted machines (gbm), a multilayer perceptron (MLP), Q-FSNet, and Q-DirichNet. All results are averaged over 500 bootstrap trials, with error bars representing 95% confidence intervals. (A) Quantile loss across target quantiles (lower values indicate better predictive accuracy). (B) Empirical coverage probability, defined as the proportion of observed responses falling below the predicted quantile, with values closer to the nominal quantile indicating better calibration (red dashed line).

Across all datasets, Q-FSNet exhibited performance comparable to or better than the baseline methods. In terms of quantile loss, Lasso and GBM achieved lower loss at lower quantiles (0.1 and 0.3). However, at higher quantiles (0.5, 0.7, and 0.9), Q-FSNet consistently outperformed Lasso and GBM. With respect to empirical coverage probability, Q-FSNet showed smaller deviations between the nominal quantile levels and the observed coverage, indicating a better calibration. We attribute this to the flexibility of the neural network architecture combined with effective noise reduction via feature selection, which allowed Q-FSNet to better capture distributional complexity in the upper tails while remaining robust to noise from irrelevant features. The Q-DirichNet model improved upon MLP but did not outperform Q-FSNet or the other baseline methods. A plausible explanation is that the Dirichlet prior induced feature weight shrinkage but weights did not reach exactly zero, limiting the model’s ability to fully discard irrelevant or noisy features.

### 2.2 Application to the Canadian Longitudinal Study on Aging Cohort

We applied our methods to the CLSA cohort to investigate potential nonlinear associations between metabolites and biological age acceleration. Although the CLSA is a longitudinal study, our analyses were restricted to baseline data, yielding a cross-sectional design, and all analyses were conducted separately for males and females. Lifestyle factors, including alcohol consumption, smoking frequency, education level, and waist-to-hip ratio, were incorporated as covariates to account for their potential confounding effects.

Prior to identifying metabolites with sweet spots, we evaluated the performance of our proposed methods on the CLSA Comprehensive cohort (CLSA-COM; [27]) and compared them with the baseline models (Lasso, GBM, and MLP) using quantile loss and empirical coverage probability as metrics, as illustrated in Figure 3. Results from Wilcoxon’s signed-rank test (Table 1) provide statistical evidence that Q-FSNet outperformed GBM and MLP in terms of mean quantile loss, and outperformed all three baseline models in the empirical coverage bias. These results were derived from pairwise comparisons of mean losses and coverage biases across all quantiles and sexes, with *p*-values adjusted using the Bonferroni method. Although Q-FSNet did not consistently outperform Lasso, it demonstrated smaller uncertainty in both quantile loss and coverage probability across the majority of cases. Additionally, Q-FSNet exhibited significantly less, and more stable training time compared to Q-DirichNet. Specifically, Q-FSNet had a mean training time of 29 minutes with a standard deviation of 5.3 minutes, while Q-DirichNet had a mean training time of 3.7 hours with a standard deviation of 1.3 hours. Because Q-DirichNet did not show statistically significant improvements over baseline methods and entailed a substantially higher computational burden, we have focused the remainder of this section on the findings obtained using Q-FSNet.

**Figure 3.**
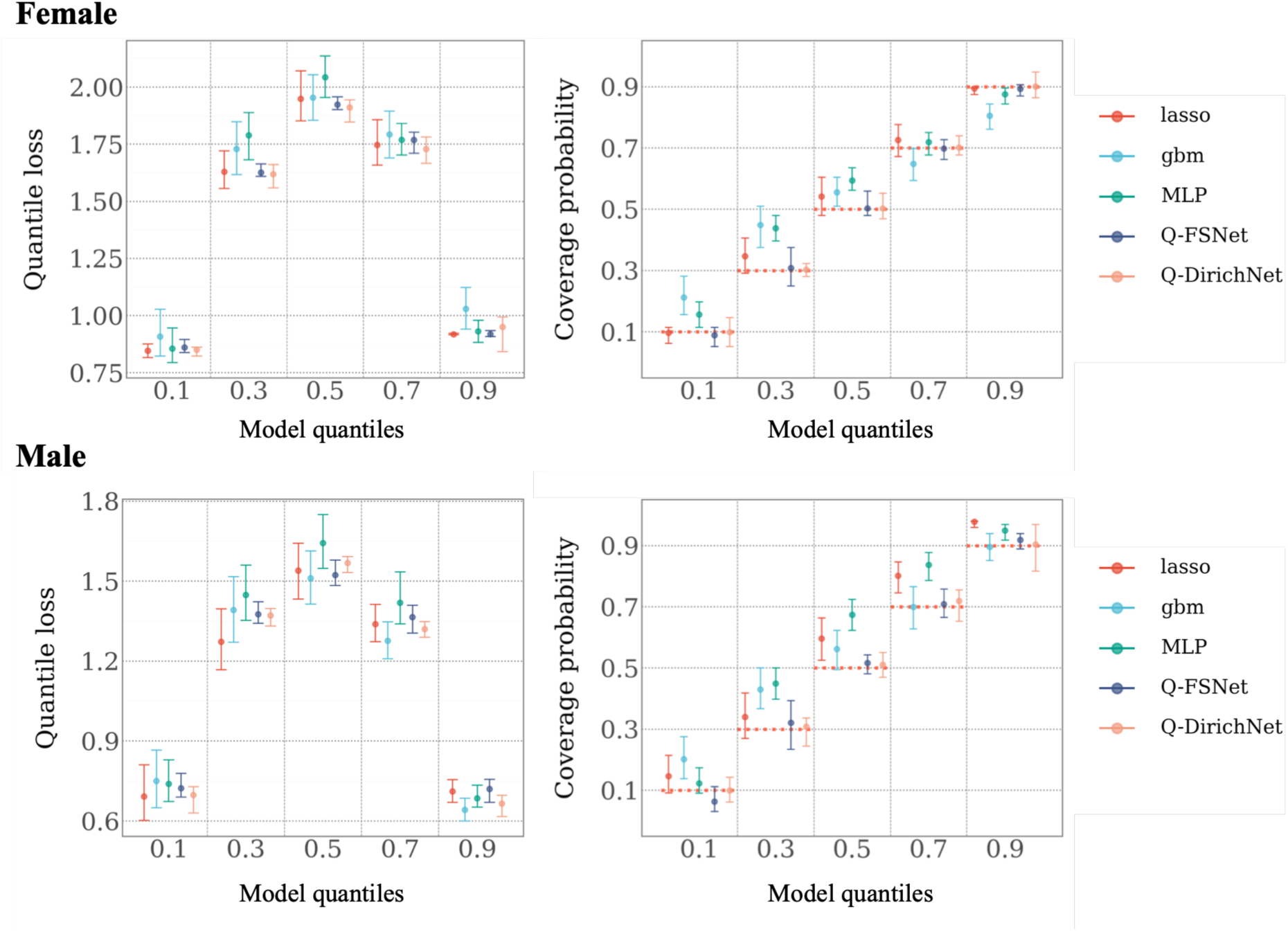
Performances of proposed models (Q-FSNet and Q-DirichNet) are compared to baseline models (lasso, gbm, and MLP) in terms of quantile losses and empirical coverage probabilities.

**Table 1.** Non-parametric hypothesis testing (Wilcoxon’s W test) results for Q-FSNet versus baseline quantile regression models (Lasso, GBM, and MLP).

| H0<br>(Quantile loss) | Adj. *p*-value<br>(Wilcoxon’s W test) | H0<br>(Coverage bias) | Adj. *p*-value<br>(Wilcoxon’s W test) |
| --- | --- | --- | --- |
| Q-FSNet > lasso | 0.988 | Q-FSNet > lasso | 0.002 |
| Q-FSNet > gbm | 0.011 | Q-FSNet > gbm | <0.0001 |
| Q-FSNet > MLP | <0.0001 | Q-FSNet > MLP | <0.0001 |

\* Coverage bias: Deviation of the empirical coverage probability from the nominal target quantiles.

#### 2.2.1 Metabolites with sweet spots and explanations

Among the total of 804 metabolites in CLSA, we focused on the 102 metabolites assigned non-zero feature weights identified by Q-FSNet. Within this subset, we discovered 25 metabolites exhibiting potential physiological sweet spots associated with biological age acceleration. To assess whether our approach captures biologically meaningful signals, we highlight six of these 25 metabolites and discuss their plausible biological interpretations, together with their partial dependence (PD) plots shown in Figure 4. The remaining metabolites and their corresponding PD plots and interpretations are provided in Figure S1 and Table S1 of the Supplementary Materials.

**Figure 4.**
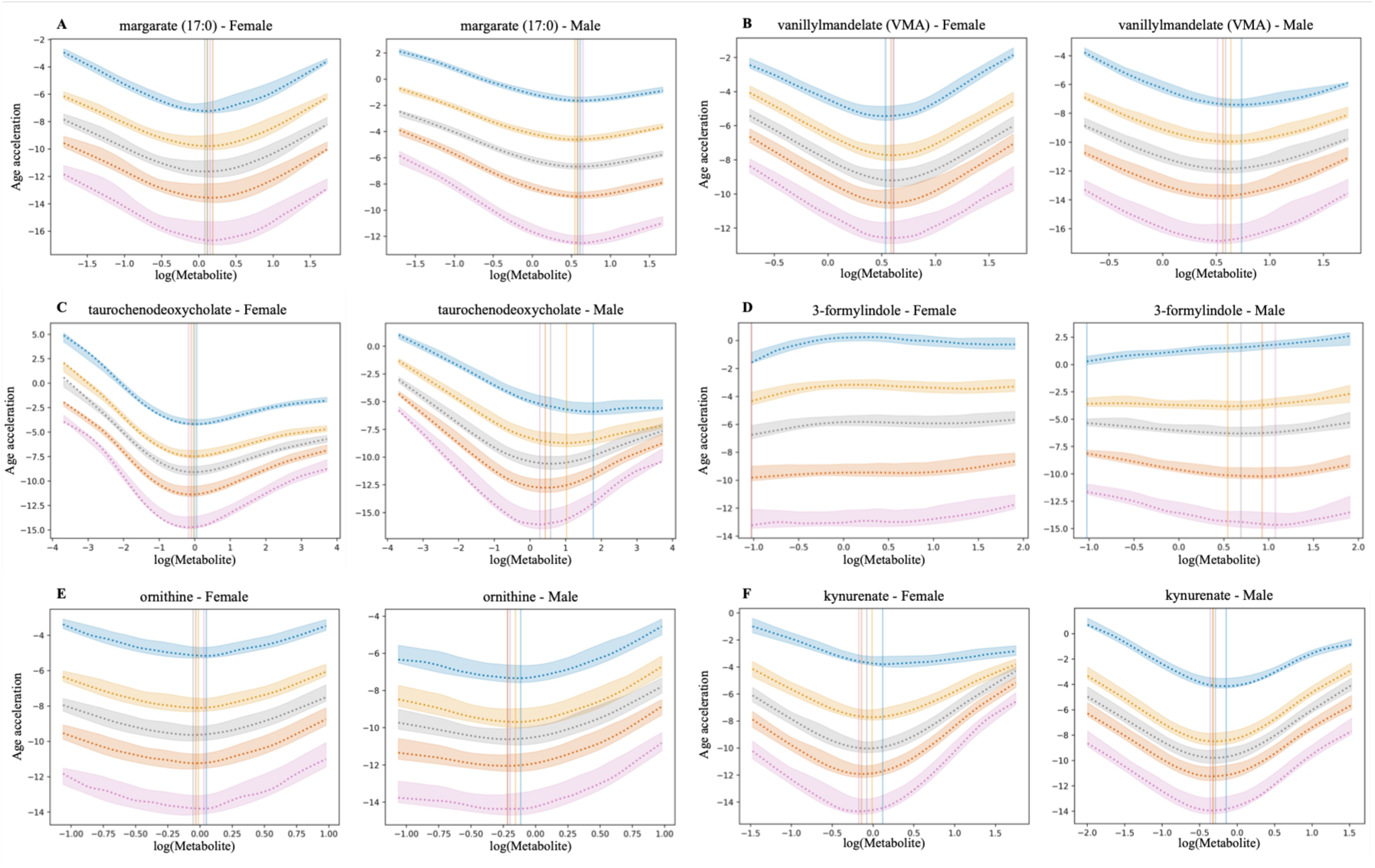
Partial dependence plots of the metabolites with optimal levels (sweet spots) discovered using Q-FSNet on the CLSA data, stratified by sex. Curves represent the quantiles of age acceleration (0.9, 0.7, 0.5, 0.3, 0.1), from top to bottom. Shaded area indicates the 95% confidence intervals obtained through MC dropouts. Vertical lines mark the sweet spots, with colour matching the shades.

**Margarate** (margaric acid) is a saturated odd-chain fatty acid occurring as trace component in ruminant milk fat and dairy-derived fats [36], and accounts for approximately 0.5% of total fatty acids in whole-fat cow milk. It participates in lipid metabolism, and can influence membrane fluidity and contribute to energy homeostasis via mitochondrial *β*-oxidation [37]. Epidemiological studies have reported inverse association between circulating margarate levels and the risk of cardiometabolic disease risk, type-2 diabetes, and systemic inflammation [37]. As shown in the partial dependence (PD) plot in Figure 4A, our results suggest that a metabolomic sweet spot, where biological age acceleration is the lowest, is present across all analyzed quantiles. Outside this range, the marginal benefit diminishes and the biological age acceleration increases. This pattern implies that maintaining margarate within this optimal range, rather than continuously increasing intake, may be important for minimizing biological aging. Notably, this sweet spot effect was more pronounced in females than males. In males, the estimated sweet spot level was higher, and the attenuation of benefit beyond the sweet spot occurred more gradually, suggesting a greater capacity to tolerate or use higher margarate levels.

**Vanillylmandelate** (vanillylmandelic acid; VMA) is a methoxyphenol and major urinary end-product of epinephrine and norepinephrine metabolism. It is formed primarily in liver through actions of catechol O-methyltransferase (COMT) on 3,4-dihydroxymandelic acid and aldehyde dehydrogenase (ALDH) on 3-methoxy-4-hydroxyphenylglycolaldehyde [38,39]. Elevated levels are associated with conditions such as chronic stress and hypertension [40]. Conversely, abnormally low levels may indicate adrenal insufficiency resulting from reduced catecholamine synthesis or the use of medications that suppress adrenal function [41]. The PD plot in Figure 4B reveals a metabolomic sweet spot across all analyzed quantiles, with similar optimal values in both females and males. This consistency suggests that the homeostatic requirement for VMA balance is largely uniform across biological aging strata and sex.

**Taurochenodeoxycholate** (Taurochenodeoxycholic acid; TCDCA) is a primary bile acid formed in the liver by conjugation of chenodeoxycholic acid with taurine through bile acid–CoA:amino acid N-acyltransferase (BAAT) [42]. It regulates cholesterol homeostasis and fat absorption. Elevated serum levels associate with drug-induced liver injury (DILI) severity and chronic hepatitis B [43], whereas low levels are associated with type 2 diabetes, obesity, and hyperlipidemia in mice [44]. We found that while the overall association is inverse, the relationship between TCDCA and biological age is not strictly linear (Figure 4C). Notably, in males the sweet spot for the 0.9 quantile was higher than the sweet spots for lower quantiles. This suggests that individuals with a higher baseline biological age may require more TCDCA to achieve optimal biological function than those with a younger biological age.

**3-Formylindole** (indole-3-carboxaldehyde; I3A) is a tryptophan-derived metabolite produced by gut microbiota, such as *Clostridium* and *Lactobacillus* [38]. As an aryl hydrocarbon receptor (AhR) agonist, it plays a vital role in mucosal immunity by stimulating interleukin-22 (IL-22) production in intestinal immune cells. 3-Formylindole is also found in various dietary sources including beans, brussels sprouts, cucumbers, cereals, and white cabbage [38]. Lower levels have been linked to increased risk of immunoglobulin A vasculitis (IgA vasculitis; IgAV) [45], whereas higher levels have been associated with rheumatic heart disease (RHD) [46]. Our study revealed sweet spots only in males with lower biological age. The PD plot in Figure 4D shows a diverging trend between the upper and lower quantiles of biological age acceleration. In the lower quantiles (0.1 and 0.3), we observed sweet spots. In contrast, males in the highest quantile (0.9) showed a monotonic increase in age acceleration as metabolite levels rose, suggesting that higher level may be detrimental for individuals with higher biological age. Among females, the trends exhibited marginally monotonic increasing trend across all levels, with no clear optimal sweet spot identified, suggesting sex-specific physiological differences.

**Ornithine** is a non-proteinogenic amino acid that plays a central role in the urea cycle. Within this pathway, the ornithine transcarbamylase (OTC) facilitates the conversion of ornithine and carbamoyl phosphate into citrulline, a critical step for ammonia detoxification and urea synthesis. Beyond the urea cycle, ornithine serves as a precursor for polyamines (putrescine, spermidine, spermine), proline, and aids in arginine regeneration for nitric oxide production [38,47]. Clinical studies have linked elevated ornithine levels to systemic sclerosis [48], hyperornithinemia [49], gyrate atrophy of choroid/retina (OAT deficiency) [50]. Conversely, low levels are typically indicative of OTC deficiency, which can lead to hyperammonemia [49]. Figure 4E shows a clear sweet spot across all age acceleration quantiles in both sexes. However, males exhibited a sharper increase in biological age when ornithine levels exceeded the estimated sweet spot, suggesting maintaining the metabolomic level within optimal range may be more critical for mitigating biological aging in males than in females.

**Kynurenate** is a quinoline carboxylic acid produced through the transamination of L-kynurenine via kynurenine aminotransferases. It serves as an endogenous antagonist of ionotropic glutamate receptors (NMDA, AMPA, kainate) and α7-nicotinic acetylcholine receptors. It is also present in various foods including chestnut and sunflower honeys, as well as in vegetables such as cauliflower, potato, and broccoli [38,51]. The clinical relevance of kynurenate is marked by its association with diverse neurological anomalies and systemic conditions [52]. Elevated levels have been positively associated with schizophrenia, epilepsy, HIV infection, amyotrophic lateral sclerosis, and cerebral malaria. Conversely, low levels are associated with neurodegenerative diseases such as Alzheimer’s, Parkinson’s, Huntington’s, and multiple sclerosis [52]. The PD plot in Figure 4F demonstrates sweet spots across all biological age quantiles and for both sexes. This result supports the idea that kynurenate acts as a critical homeostatic regulator, where both deficiency and excess contribute to accelerated biological aging, highlighting the importance of maintaining kynurenate within a specific physiological window.

Other metabolites that we discovered to have sweet spots include 4 amino acids (1-methylhistidine, N,N,N-trimethyl-alanylproline betaine (TMAP), N-acetyltaurine, trans-urocanate), 12 lipids (1,2-dilinoleoyl-GPC (18:2/18:2), 1-(1-enyl-palmitoyl)-2-palmitoleoyl-GPC (P-16:0/16:1)*, 1-(1-enyl-stearoyl)-2-arachidonoyl-GPE (P-18:0/20:4)*, 1-oleoyl-2-arachidonoyl-GPE (18:1/20:4)*, 1-stearoyl-2-oleoyl-GPC (18:0/18:1), 4-hydroxy-2-oxoglutaric acid, glycolithocholate sulfate*, glycoursodeoxycholate, maleate, oleoylcarnitine (C18:1), adipoylcarnitine (C6-DC), 1-stearoyl-2-linoleoyl-GPE (18:0/18:2)), 1 vitamin (pyridoxate) and 2 xenobiotics (erythritol, stachydrine). PD plots and corresponding biological interpretations for these metabolites are provided in Figure S1 and Table S1 of the Supplementary Materials. Additional biological validation or complementary approaches, such as causal inference analyses, will be required to confirm and further substantiate these observations.

## 3 Discussion

This study introduces a scalable quantile regression neural network framework designed to identify physiological sweet spots within high-dimensional data. The proposed approach facilitates the smooth, data-driven detection of non-linear effects, integrates automated feature selection with uncertainty estimation, and reveals quantile-specific signatures across the outcome distribution. The selection of a neural network architecture offers distinct advantages over traditional ensemble or regression-based methods. Unlike linear regression or gradient-boosting machines, which often produce discontinuous, axis-aligned decision boundaries, neural networks learn a continuous non-linear manifold. This architectural smoothness is essential for identifying stable physiological sweet spots; it prevents the introduction of step-function artifacts and ensures that the identified optima reflect underlying biological gradients rather than local data-splitting heuristics. Q-FSNet enhances this architecture with a specialized feature selection layer and Monte Carlo (MC) dropout, effectively inducing sparsity to eliminate noisy, irrelevant features while providing quantifiable prediction uncertainties.

Applying our method to the CLSA Comprehensive dataset, we identified 25 metabolites with potential sweet spots, many of which are corroborated by their established biological functions. However, while our framework provides a robust data-driven approach for discovering non-linear associations, our results identify associations, which cannot be assumed to be causal. Our method identifies high-priority candidates for subsequent biological experimentation to assess potential causal mechanisms.

The current implementation of Q-FSNet is an extension of a standard MLP. This architectural simplicity was intentional, as more complex tabular neural network structures often require significantly larger datasets to prevent overfitting. Nevertheless, the modular nature of neural networks allows for the seamless integration of sophisticated components, such as Transformers, into this framework.

One limitation of the current approach is that the prediction uncertainties provided by MC dropout primarily account for epistemic uncertainty. The exclusion of aleatoric uncertainty may result in underestimated total uncertainty compared to fully Bayesian neural networks, such as the previously introduced Q-DirichNet. However, given that variational inference-based Bayesian networks entail a substantially higher computational burden, Q-FSNet offers a pragmatic trade-off between calibration and efficiency. Future work may involve exploring more efficient extensions of Q-DirichNet to bridge this gap. Furthermore, addressing the hierarchical and correlated nature of metabolites through grouped or adaptive Lasso-style feature selection layers may improve mechanistic insight. Lastly, while this study utilized the baseline CLSA dataset in a cross-sectional design, future research will extend this methodology to a longitudinal framework. Leveraging follow-up data from the CLSA cohort will allow us to model temporal trajectories of metabolites and assess how sweet spot stability may shift over time, providing a more dynamic understanding of metabolic health and biological aging.

This study presented Q-FSNet, a scalable quantile regression framework designed to navigate the complexities of high-dimensional data. By leveraging the continuous manifold learning of neural networks, we successfully identified 25 metabolites with physiological sweet spots associated with biological aging in the CLSA cohort, and these findings are supported by existing biological literature. Unlike traditional methods, Q-FSNet provides a smooth, data-driven alternative that integrates feature selection and uncertainty estimation without requiring a priori assumptions about non-linear relationships. While current limitations regarding aleatoric uncertainty and causal interpretation remain, the modularity of this framework provides a robust foundation for future integration with causal inference and more complex architectures. Ultimately, Q-FSNet offers a powerful tool for researchers to move beyond linear associations, uncovering the nuanced, non-linear metabolic signatures that define human health and aging.

The 25 metabolites discovered to have sweet spots relate to diverse biological processes. These include neuro-metabolic and signaling pathways (kynurenate, vanillylmandelate), muscle metabolism and amino acid turnover (1-methylhistidine, N,N,N-trimethyl-alanylproline betaine, N-acetyltaurine, trans-urocanate), and nitrogen disposal via the urea cycle (ornithine), illustrating the broad applicability of this statistical approach to elucidating physiological mechanisms. Several of these metabolites are known to relate to diet (margarate, pyridoxate, erythritol, stachydrine) or microbial-host co-metabolism (3-formylindole, taurochenodeoxycholate, glycolithocholate sulfate, and glycoursodeoxycholate). Notably, compounds like 1-methylhistidine and kynurenate reflect the intersection of endogenous turnover and dietary supply. These observations suggest that clarifying optimal intake of certain nutrients and the nature of the microbiome could provide important information for public health and the optimization of healthy aging.

## 4 Methods

This study employs a deep learning framework, the quantile feature selection network (Q-FSNet), and its Bayesian extension (Q-DirichNet), to model risk-population-specific biomarkers for ageing-related phenotypes. In this Section, we describe these methods and provide an overview of the Comprehensive Cohort of the Canadian Longitudinal Study on Aging (CLSA; [27]).

### 4.1 Quantile regression

Traditional ordinary least squares (OLS) regression models the conditional mean of the outcome *Y* given explanatory features *X*, 𝔼[*Y*|*X*]. In contrast, quantile regression (QR), first introduced by Koenker and Basset [11], models the conditional quantile *Q*_𝜏_(*Y*|*X*) at a specific quantile level 𝜏 ∈ (0,1). This allows the impact of features to be examined at the distributional tails, capturing heterogeneity and remaining robust to non-Gaussian error distributions and outliers, conditions which are common in biological data.

The objective of QR is to minimize the sum of asymmetrically weighted absolute errors:

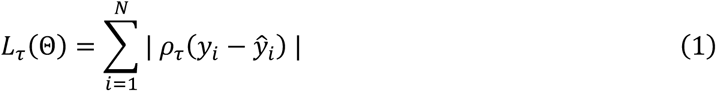

Here Θ represents the networks’ weights and biases, *ȳ_i_* represents the predicted conditional quantile for the *i*-th individual, and 𝜌_𝜏_(*u*) is the pinball loss:

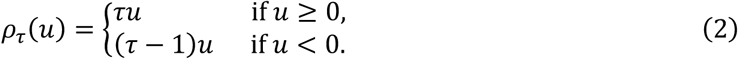

For an extreme quantile like 𝜏 = 0.9, the loss function heavily penalizes negative residuals (underestimation), driving the prediction *Ȳ* = (*ȳ*_1_, …, *ȳ*_*N*_) toward the 90^th^ percentile of the true outcome distribution.

### 4.2 The quantile regression feature selection neural network (Q-FSNet)

The quantile feature selection network (Q-FSNet) is a hybrid architecture that builds upon a standard multilayer perceptron (MLP). This architecture offers the necessary capacity to model non-linear associations between metabolome and phenotypes while being less prone to overfitting compared to more complex architectures. While deep neural networks with many hidden layers or more sophisticated architectures like the attention mechanism are available [16], they require a substantial amount of data and are susceptible to overfitting when the data size is limited. We illustrate the main structure of Q-FSNet in Figure 5.

**Figure 5.**
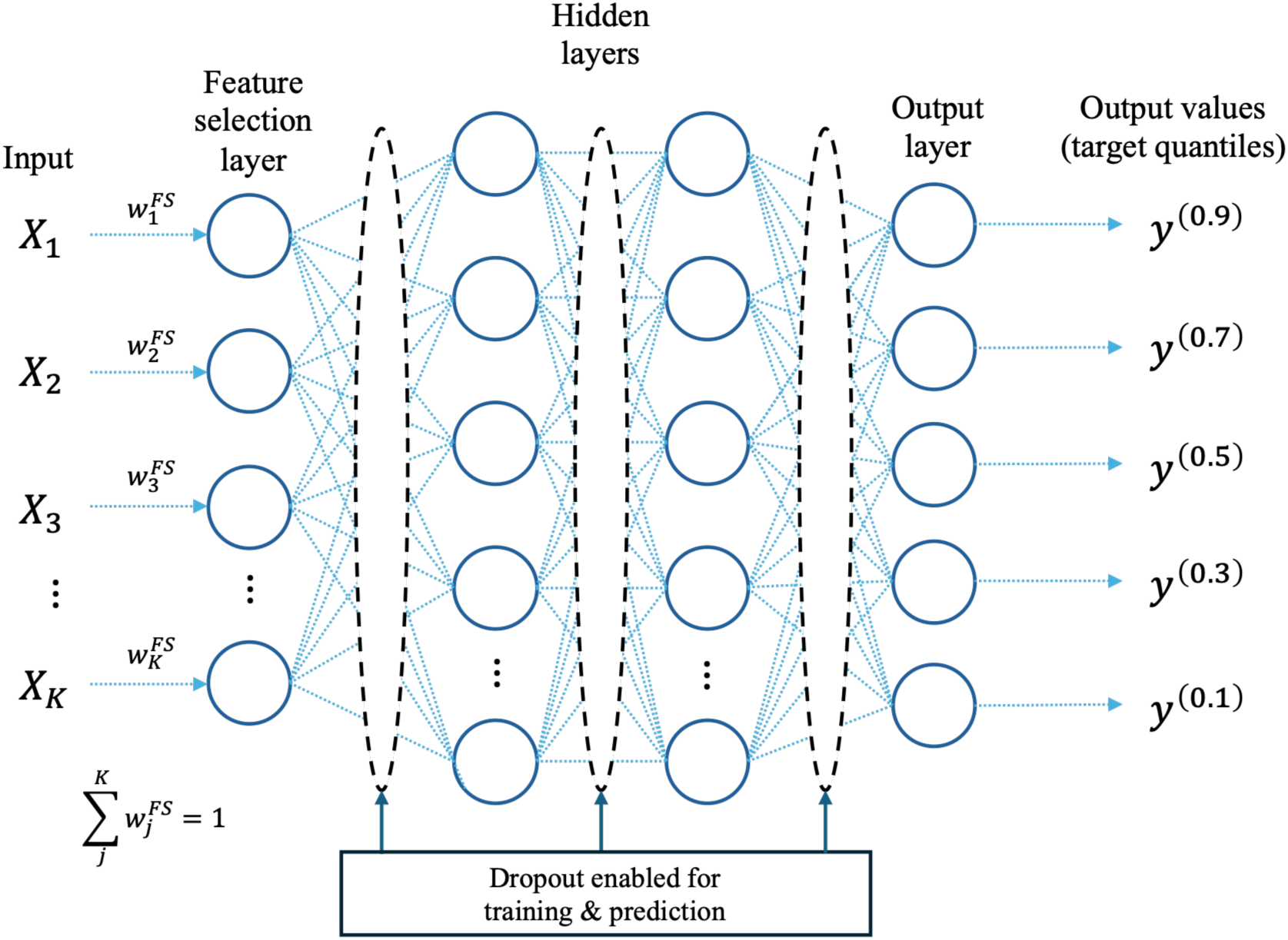
The architecture of Q-FSNet. Feature selection layer is situated between the input data and the hidden layers, utilizing one-to-one connections to the input features. The learned feature weights control the signal flow into the network, serving as a direct measure of feature importance. To enforce sparsity and promote competitive weighting, these weights are constrained to sum to one. During inference, Monte Carlo (MC) dropout is applied within the hidden layers to approximate Bayesian uncertainty estimation.

#### 4.2.1 Feature selection layer

To counter the susceptibility of NNs to uninformative features in high-dimensional tabular settings [16,19,20], we utilize a feature selection (FS) layer [25], a technique embedded directly within the NN. The FS layer is connected immediately after the input layer. It consists of a vector of trainable weights *W*^*FS*^, wherein each weight 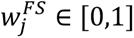 corresponds to the importance of the *j*-th input feature *X*_j_where *j* ∈ {1, …, *K*}. The output of the FS layer for an input row *x*_*i*_ is the element-wise multiplication:

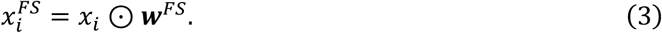

Here ⊙ denotes the Hadamard product. To enforce sparsity and promote a unique solution for which weights of irrelevant features converge to zero, the training of *W*^*FS*^ is coupled with an *L*_1_-penalty, and a constraint that encourages the weights to sum to one:

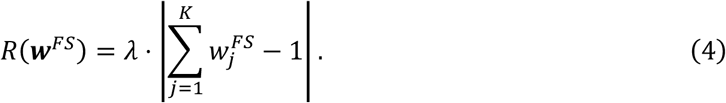

Here *K* is the number of input features and 𝜆 controls the regularization strength. This constraint ensures that the importance of each feature is influenced by the interaction with other features, effectively allowing the network to perform feature selection in an end-to-end manner.

#### 4.2.2 Prediction uncertainty estimation using Monte Carlo dropout

To address the lack of prediction uncertainty in deterministic NNs, we employ Monte Carlo dropout [26]. Dropout, originally a regularization technique, is typically active only during training. By leaving dropout active during the prediction phase, the model can approximate a Bayesian neural network [26,53]. Running *T* forward passes for a single input data *X*, each with a different set of dropout connections, yields a distribution of predictions *Ȳ*^(*t*)^ for *t* = 1, …, *T*. The average of these predictions serves as the final prediction. This method provides a computationally efficient and tractable means of quantifying model trustworthiness.

### 4.3 Bayesian extension to Q-FSNet (Q-DirichNet)

To provide a fully Bayesian extension of the feature selection layer process, we introduce Quantile Dirichlet network (Q-DirichNet). In this extension, the deterministic neural network parameters Θ (including the feature selection weights *W*^*FS*^) are replaced by random variables within a hierarchical Bayesian framework. Consequently, Q-DirichNet learns its parameters and priors directly from the data, retaining Q-FSNet’s feature selection capability without requiring a regularization parameter, and all while offering the inherent advantage of uncertainty estimation.

#### 4.3.1 Dirichlet prior for feature selection

We place a Dirichlet prior *P*(*W*^*FS*^) over the feature selection layer weights. This Dirichlet distribution Dir(α), naturally constrains the FS layer weights to be non-negative and sum to one:

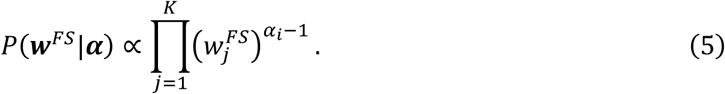

Here α = (α_1_, …, a_*K*_) is a vector of the concentration parameters, which are themselves random variables α_j_ ∼ Gamma(1,1), and learned from the data. This structure enforces sparsity and feature interaction constraint without relying on the external *L*_1_ regularization used in Q-FSNet. Specifically, lower learned values of α_*i*_ indicates that the model has identified the *j*-th feature as less relevant, pushing it corresponding weight 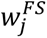 toward zero.

#### 4.3.2 Asymmetric Laplace distribution likelihood

To establish a Bayesian framework compatible with QR, we model the observations *Y* using an asymmetric Laplace distribution (ALD). The negative log-likelihood of this distribution is equivalent to the pinball loss [54,55]. For a given quantile 𝜏, the likelihood is defined as:

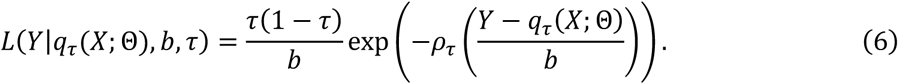

Here *q*_𝜏_(*X*; Θ) is the network’s prediction for the 𝜏-th quantile, *b* is the scale parameter related to the ALD’s spread, and 𝜌_𝜏_(⋅) represents the pinball loss. In the hierarchical design, *b* is also modeled as a random variable *b* ∼ HalfCauchy(*β*_*b*_), the probability density function of which with its scale parameter *β*_*b*_ is defined as:

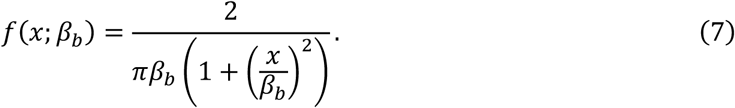

The half-Cauchy distribution is a common choice for scale parameters due to its non-negative support and heavy tails, which create a weakly informative prior [56–58]. To further stabilize the model, we applied a half-Normal hyperprior to *β*_*b*_ ∼ HalfNormal(0, 10). This constraint prevents *β*_*b*_from collapsing toward zero, thereby mitigating computational instability while remaining highly flexible. This hierarchical approach provides a more robust fit by learning the level of observation noise and prior structure directly from the data. Finally, the posterior distributions for all parameters are then estimated using Markov chain Monte Carlo (MCMC) methods.

### 4.4 Sweet spot identification

To identify population-level metabolomic optima, we estimated marginalized response functions for each metabolite *X*_j_. Specifically, we calculated the partial dependence (PD) of the outcome phenotype value on each metabolite *X*_j_ at value *x*_j_ as follows:

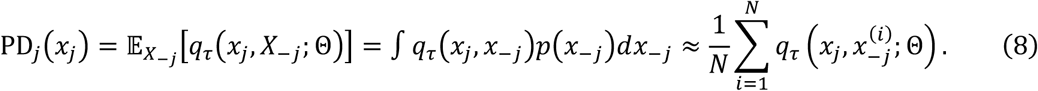

Here *p*(*x*_–j_) is the marginal density function of the non-target features *X*_–j_, and *x*^(*i*)^ represents the observed values of all other metabolites for individual *i*. This marginalization ensures that the identified trends are representative of the average population response rather than being driven by specific individual metabolic contexts. For each metabolite, PD was evaluated over its observed values. Then, the *potential sweet spot* is defined as the metabolite level that minimizes the marginalized prediction within the observed biological range:

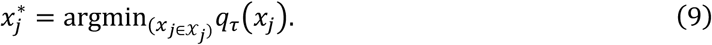

Here *Y*_j_ is the observed range of the metabolite *X*_j_. To ensure the robustness of these optima and distinguish them from strictly monotonic trends, we applied a non-monotonicity filter. A metabolite was only categorized as having a sweet spot only if the estimated 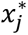 lay within the central portion of the observed distribution (e.g., between the 25^th^ and 75^th^ percentiles). This criterion reduces the risk of misidentification of boundary effects where the model may show minimums at extreme, sparsely populated concentrations, as biologically meaningful homeostatic set-points.

### 4.5 The Canadian Longitudinal Study on Aging cohort

Our selected subset for analysis from the Canadian Longitudinal Study on Aging Comprehensive Cohort (CLSA-COM; [27]) comprised 30,097 participants aged between 45 and 85 years. Of these participants, 52.3% were female, and the average age was 63.1 years at time of the blood draw for metabolomic measurements. 94% of participants self-reported as belonging to the White ethnic group, followed by Asian (2%), and Black (0.7%). Their metabolites and lifestyle factors (alcohol consumption, smoking frequency, education level, and waist-hip ratio) were included as the input features. Biological age acceleration is defined as the difference between the participants’ absolute epigenetic age estimates and chronological age. These measures were provided by the CLSA as part of the released dataset.

#### 4.5.1 Quality control and data preprocessing

From a total of 1,458 normalized metabolites, 295 metabolites whose super pathways are unknown were excluded. Then, an additional 359 metabolites with more than 10% missing values were further excluded, resulting in a total of 804 metabolites. Remaining missing values were imputed using the multiple imputation method implemented in the *mice* package in R (version 4.4.2). The normalized metabolite levels were further log-transformed and standardized prior to model training to address data skewness and improve model training stability. At the participant level, 1,288 individuals with non-missing biological age acceleration (635 males, 653 females) were included in the analyses. Among the 804 metabolites, Q-FSNet assigned non-zero feature weights to 102 metabolites. Examining the partial dependence of biological acceleration on these metabolites further identified 25 metabolites exhibiting physiological sweet spots.

### 4.6 Model training

Data were first split into training and test (held-out) sets, and the training set is further split into training and validation sets. The training, validation, test set ratio was 7:1.5:1.5. Hyperparameter tunings were carried out using the training and the validation sets, and final metrics of the best models were calculated using the held-out set.

Three baseline quantile regression models were employed for comparative analysis, including Lasso-regularized linear regression (lasso; [59,60]), gradient boosted machine (gbm; [13,61]), and multi layer perceptron (MLP; [14,15]). The lasso models’ regularization parameter was tuned via 5-fold cross validation. The hyperparameters of the gbm and NN-based models (MLP, Q-FSNet, Q-DirichNet) models were tuned using Bayesian optimization [62,63]. For the NN-based methods, AdamW optimizer [64,65] and GeLU [66] activation functions were used.

### 4.7 Software

The following modules were employed in this project under Python (version 3.10.13; [67] environment: *PyTorch* (version 2.7.1; [68]) is used for training Q-FSNet, *Pyro* (version 1.9.1; [69]) is used for training Q-DirichNet. The python packages *statsmodels* [70] is used for Lasso quantile regression, and *lightgbm* [71] for gbm-based quantile regression. Hyperparameter tuning using Bayesian optimization was carried out using the *ray* [72] module. For data preprocessing and imputation, R-software (version 4.4.2; [73]) is used, along with the packages *tidyverse* [74] and *mice* [75].

## 5 Acknowledgements

This research was made possible using the data/biospecimens collected by the Canadian Longitudinal Study on Aging (CLSA). Funding for the Canadian Longitudinal Study on Aging (CLSA) is provided by the Government of Canada through the Canadian Institutes of Health Research (CIHR) under grant reference: LSA 94473 and the Canada Foundation for Innovation, as well as the following provinces, Newfoundland, Nova Scotia, Quebec, Ontario, Manitoba, Alberta, and British Columbia. This research has been conducted using the following CLSA datasets: the CLSA Baseline Comprehensive Dataset version 7.0, Epigenetics data version 1.1, and Metabolomics data version 2.0 under Application Number 2206033. The CLSA is led by Drs. Parminder Raina, Christina Wolfson and Susan Kirkland. The time and commitment of the participants to the CLSA study platform is gratefully acknowledged, without whom this research would not be possible.

The opinions expressed in this manuscript are the author’s own and do not necessarily reflect the views of the Canadian Longitudinal Study on Aging.

## 6 Authors’ contributions

J.M. was involved in the conceptualization, data curation, visualization, and methodology of the study, also contributed with software used, conducted investigation, formal analysis, wrote, reviewed, and edited the original draft. O.V. was involved in the data curation, writing, review, and editing. A.B.-W. was involved in writing, review, editing, supervision, and project administration. L.E. was involved in writing, review, editing, supervision, and project administration.

## 8 Data availability

Data are available from the Canadian Longitudinal Study on Aging (www.clsa-elcv.ca) for researchers who meet the criteria for access to de-identified CLSA data.

