## Supplementary Materials for "A New Sparse Bayesian Quantile Neural Network-based Approach and Its Application to Discover Physiological Sweet Spots in the Canadian Longitudinal Study on Aging"

**Figure S1.** Partial dependence plots of the metabolites with sweet spots related to biological aging

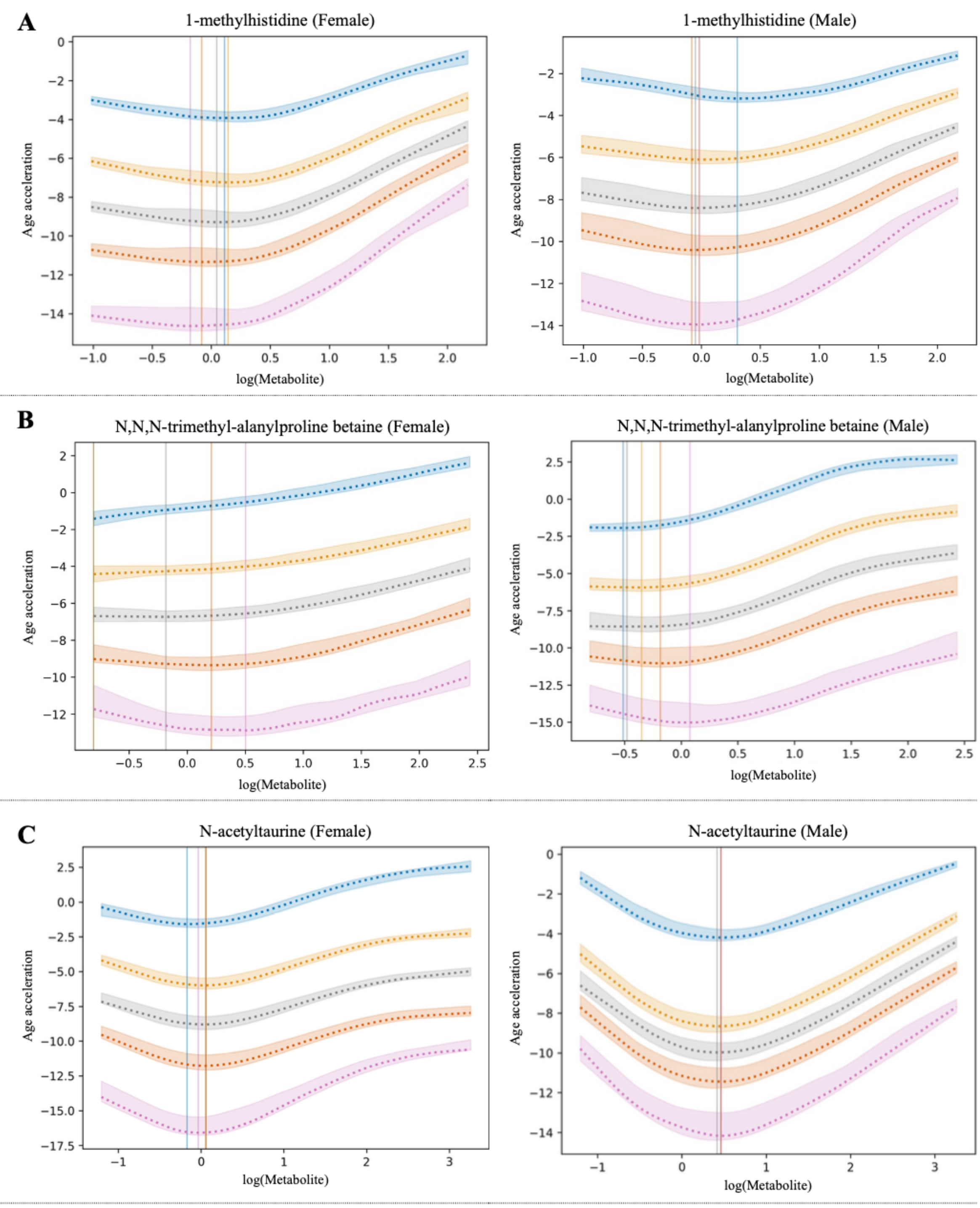

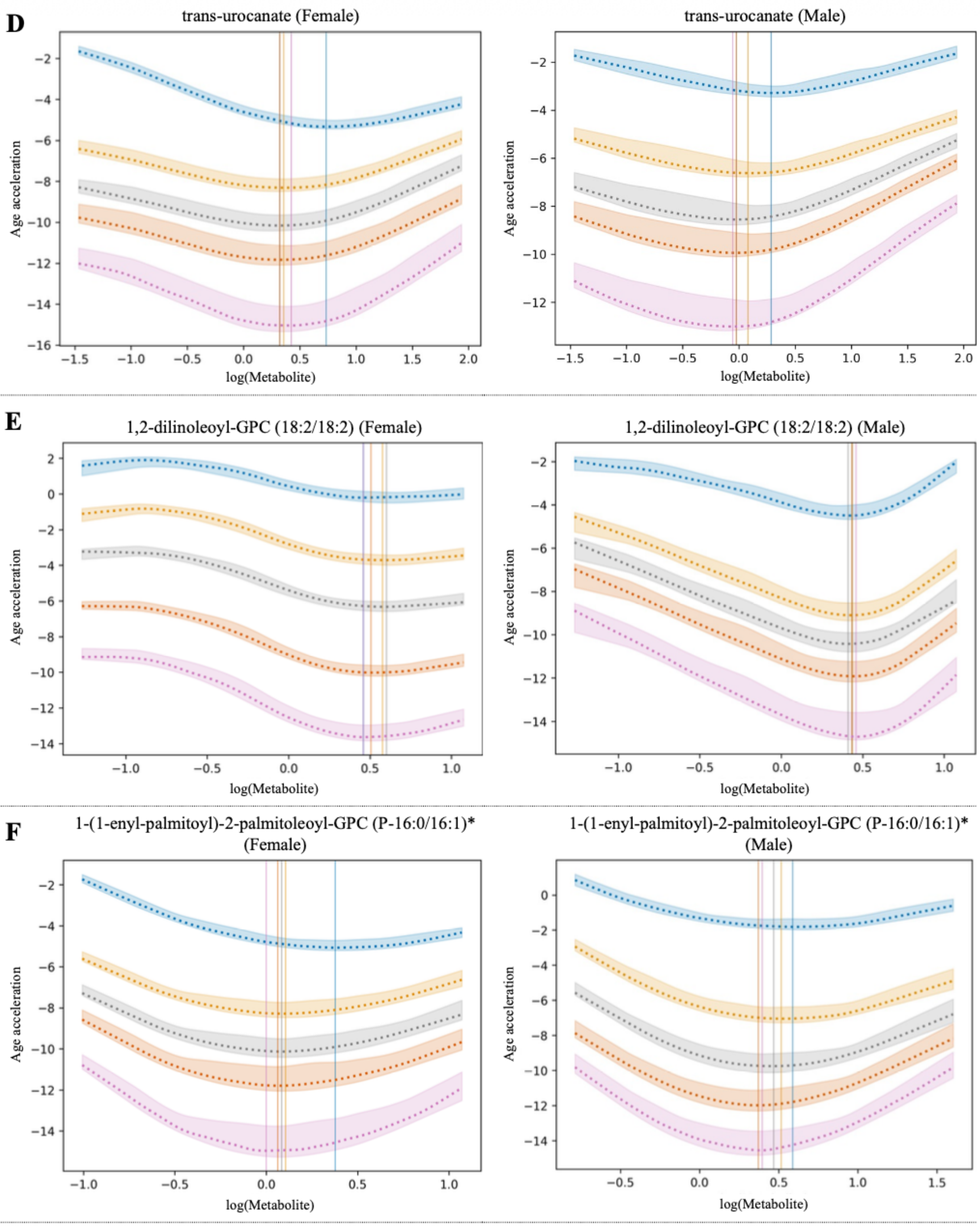
D
trans-urocanate (Female)
trans-urocanate (Male)
Age acceleration
log(Metabolite)
E
1,2-dilinoleoyl-GPC (18:2/18:2) (Female)
1,2-dilinoleoyl-GPC (18:2/18:2) (Male)
Age acceleration
log(Metabolite)
F
1-(1-enyl-palmitoyl)-2-palmitoleoyl-GPC (P-16:0/16:1)* (Female)
1-(1-enyl-palmitoyl)-2-palmitoleoyl-GPC (P-16:0/16:1)* (Male)
Age acceleration
log(Metabolite)

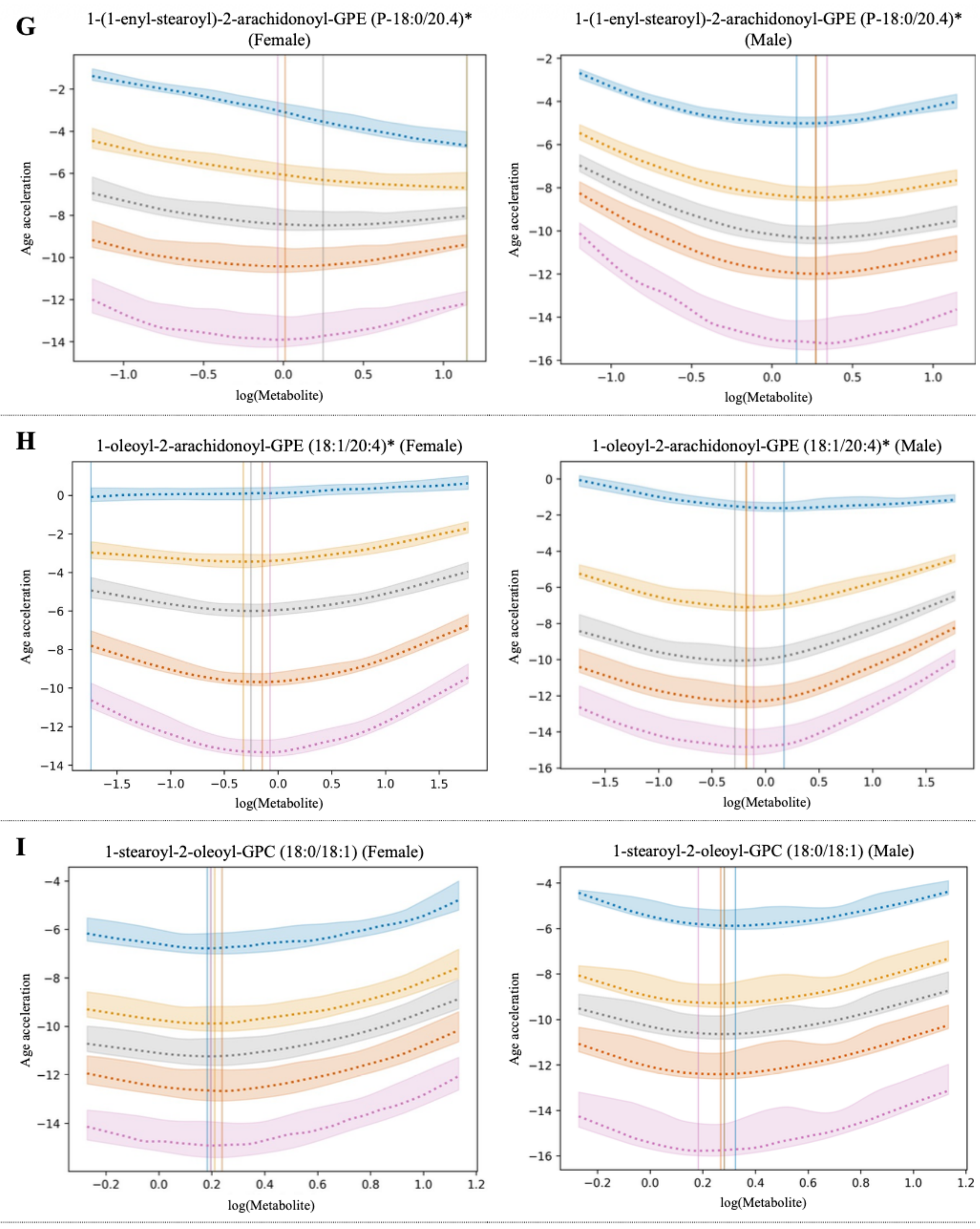
G
1-(1-enyl-stearoyl)-2-arachidonoyl-GPE (P-18:0/20.4)* (Female)
1-(1-enyl-stearoyl)-2-arachidonoyl-GPE (P-18:0/20.4)* (Male)
H
1-oleoyl-2-arachidonoyl-GPE (18:1/20:4)* (Female)
1-oleoyl-2-arachidonoyl-GPE (18:1/20:4)* (Male)
I
1-stearoyl-2-oleoyl-GPC (18:0/18:1) (Female)
1-stearoyl-2-oleoyl-GPC (18:0/18:1) (Male)
Age acceleration
log(Metabolite)

**J**

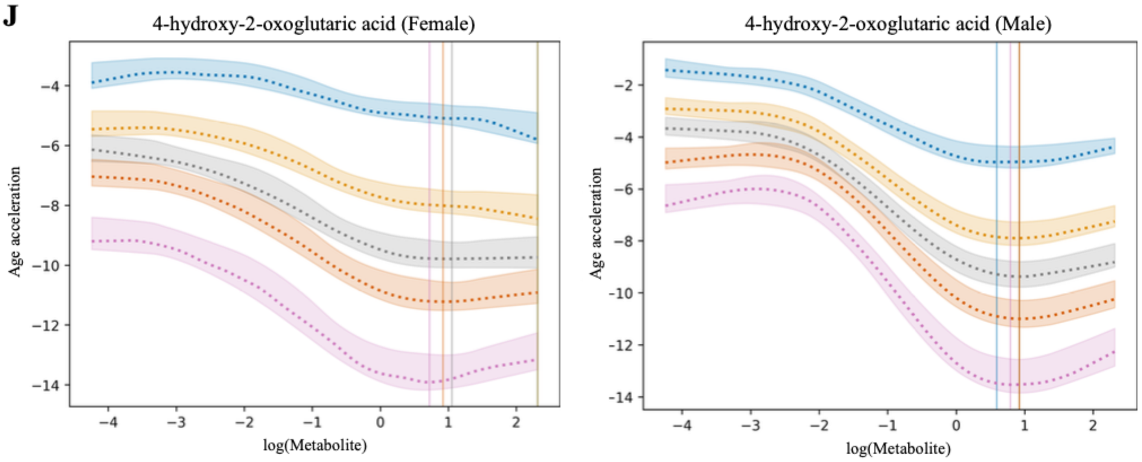

4-hydroxy-2-oxoglutaric acid (Female)
Age acceleration
log(Metabolite)
4-hydroxy-2-oxoglutaric acid (Male)
Age acceleration
log(Metabolite)


**K**

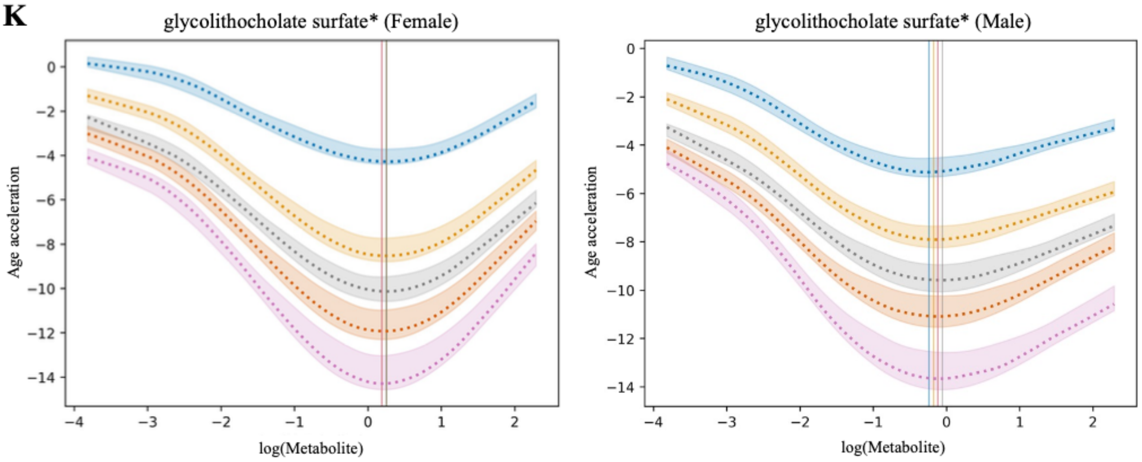

glycolithocholate surfate* (Female)
Age acceleration
log(Metabolite)
glycolithocholate surfate* (Male)
Age acceleration
log(Metabolite)


**L**

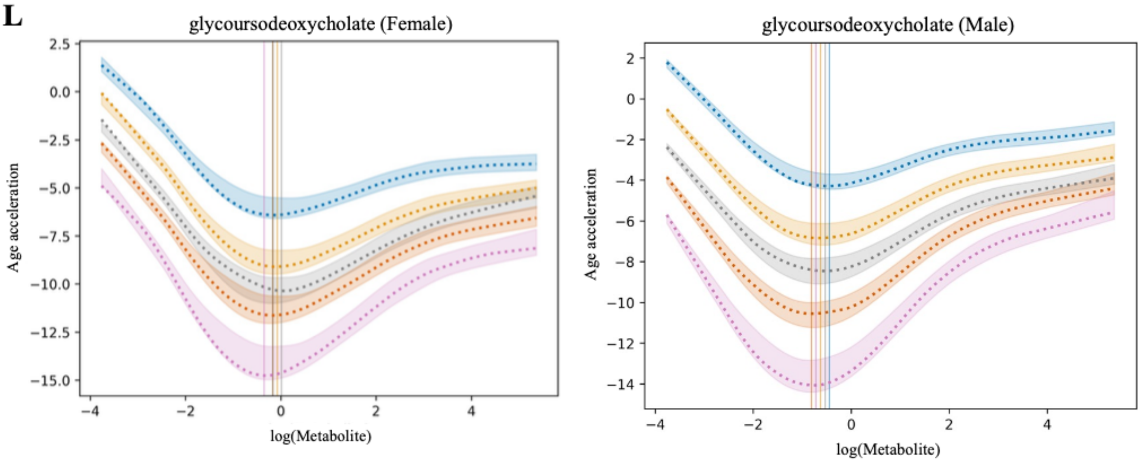

glycoursodeoxycholate (Female)
Age acceleration
log(Metabolite)
glycoursodeoxycholate (Male)
Age acceleration
log(Metabolite)

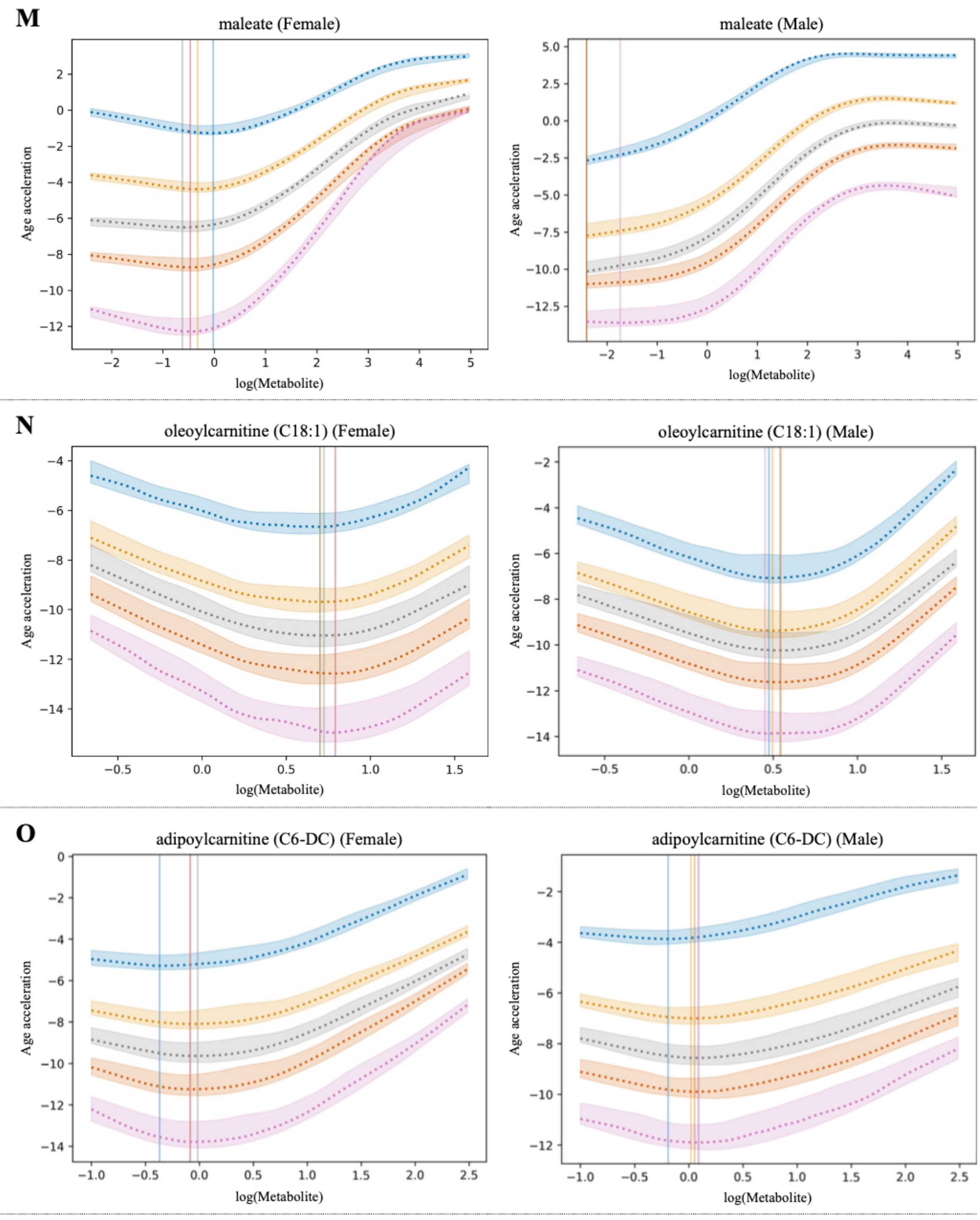
M
maleate (Female)
maleate (Male)
N
oleoylcarnitine (C18:1) (Female)
oleoylcarnitine (C18:1) (Male)
O
adipoylcarnitine (C6-DC) (Female)
adipoylcarnitine (C6-DC) (Male)
Age acceleration
log(Metabolite)

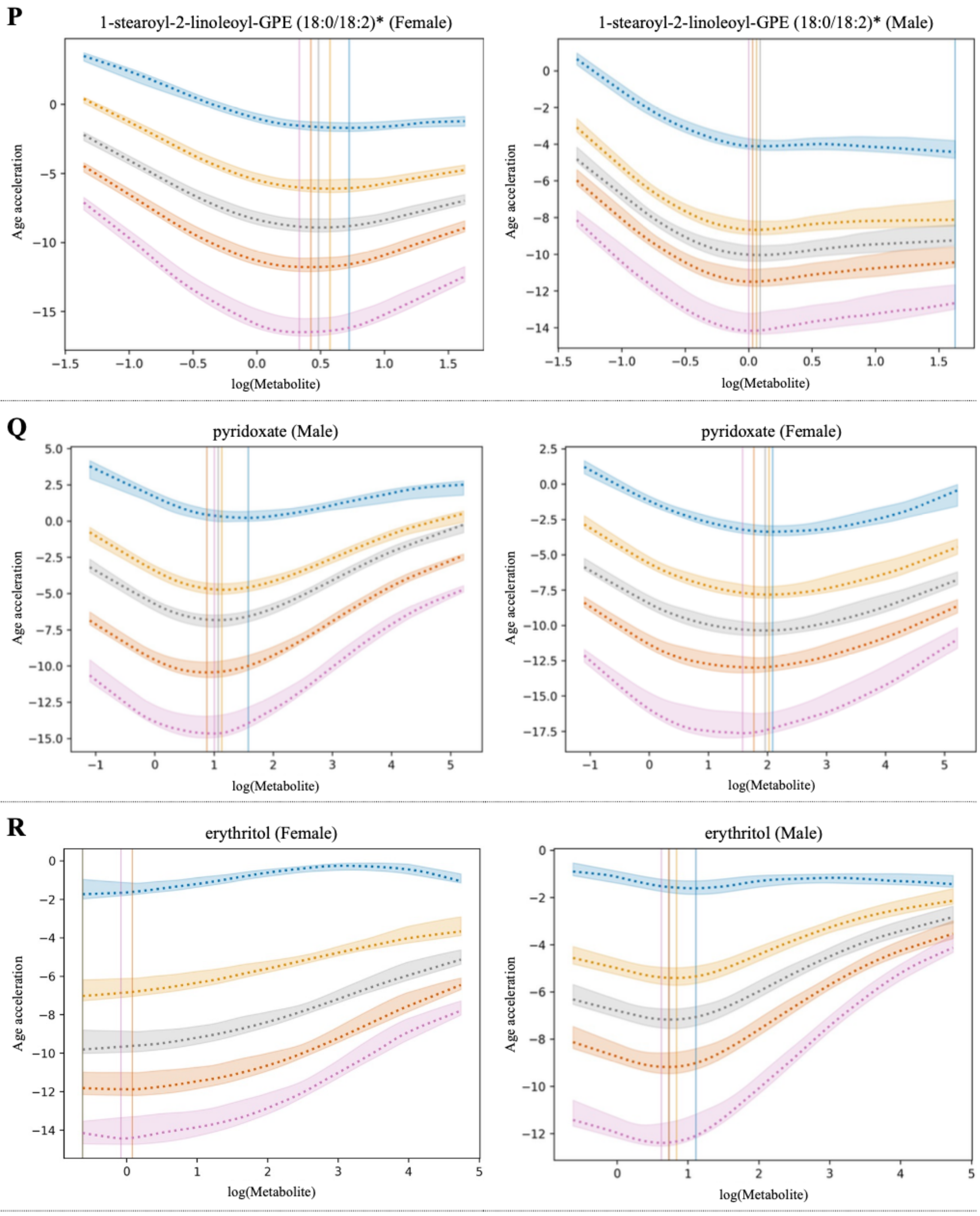
P
1-stearoyl-2-linoleoyl-GPE (18:0/18:2)* (Female)
1-stearoyl-2-linoleoyl-GPE (18:0/18:2)* (Male)
Q
pyridoxate (Male)
pyridoxate (Female)
R
erythritol (Female)
erythritol (Male)
Age acceleration
log(Metabolite)

**S**

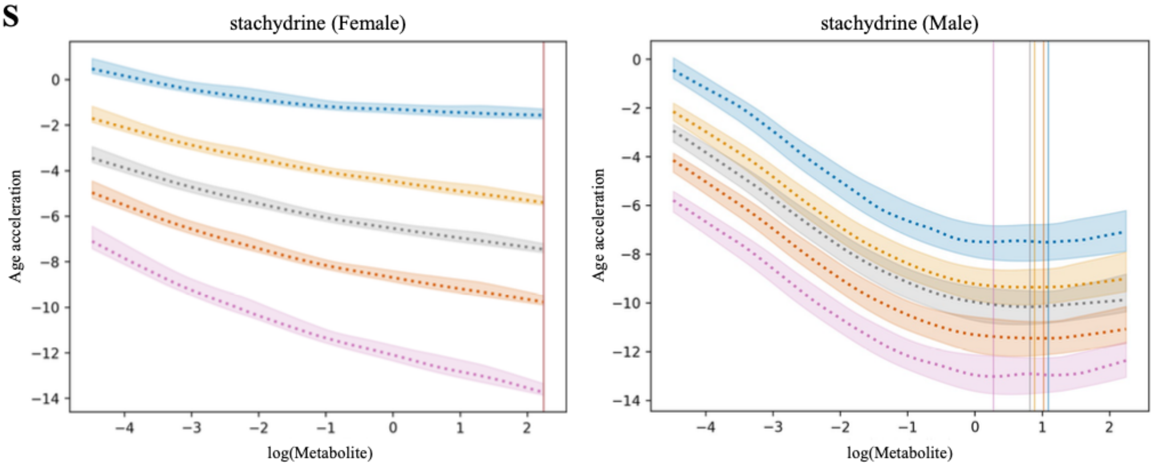
stachydrine (Female)
stachydrine (Male)
Age acceleration
log(Metabolite)
0
−2
−4
−6
−8
−10
−12
−14
−4
−3
−2
−1
0
1
2

**Table S1.** Additional metabolites exhibiting physiological sweet spots associated with biological aging identified by Q-FSNet in CLSA data. Citations correspond to the reference list in the main text.

| Fig | Metabolite (HMDB) | Super pathway | Roles/related studies/identified associations |
|---|---|---|---|
| S1A | 1-methylhistidine (HMDB0000001) | Amino Acid | • An amino acid derivative and dietary biomarker that largely reflects intake of anserine-rich meats (especially poultry and other muscle foods), rather than endogenous muscle catabolism, with higher plasma and urinary levels consistently observed in omnivores compared with vegetarians. It has been used as an objective indicator of habitual meat consumption in nutritional and epidemiologic studies [38,76].<br>• Elevated level associates with increased risk of several conditions and diseases, including acute myocardial infarction [77], hypertension [78,79], systemic sclerosis with joint pain [48], Alzheimer's [80], preeclampsia [81], and chronic kidney disease [82].<br>• Sweet spots were found across all biological age acceleration quantiles in both sexes. In the context of its dietary origin, very low 1-methylhistidine concentrations may indicate little or no meat intake and potentially reflect low intake of meat-derived nutrients, whereas very high concentrations may indicate dietary patterns or comorbid states linked to higher cardiometabolic and kidney disease risk. |
| S1B | N,N,N-trimethyl-alanylproline betaine (TMAP) (HMDB0240365) | Amino Acid | • A dipeptide betaine consisting of N,N,N-trimethyl-L-alanine linked to L-proline via a peptide bond. It may derive from myosin light chain protein degradation [38,83].<br>• Elevated levels associate with reduced kidney function, including CKD [83,84].<br>• In males, physiological sweet spots were consistently identified at low concentrations across all quantiles. Conversely, females only exhibited these sweet spots within the lower quantiles (0.1–0.5). Within the higher female quantiles, the relationship shifted to a monotonically increasing pattern, aligning with literature that associates higher TMAP levels with systemic decline. These findings suggest that while TMAP generally correlates positively with aging, its impact is non-linear and dependent on an individual's baseline aging rate, providing a quantile-specific therapeutic window to preserve their physiological resilience. |
| S1C | N-acetyltaurine (HMDB0240253) | Amino Acid | • A highly water-soluble organosulfonic acid formed by N-acetylation of taurine with acetate, catalyzed by NAT synthase. It is an endogenous metabolite and a biomarker of ethanol metabolism and endurance exercise [38,85].<br>• Elevated levels associate with hyperacetatemia [86], while low levels are linked to higher abdominal obesity, hypertension, inflammation, and prevalence of type 2 diabetes [87].<br>• Physiological sweet spots were identified across all biological age acceleration quantiles for both sexes, aligning with the dual-risk nature of the metabolite reported in previous research. |

| Fig | Metabolite (HMDB) | Super pathway | Roles/related studies/identified associations |
|---|---|---|---|
| S1D | trans-urocanate (HMDB0000301) | Amino Acid | • A α,β-unsaturated monocarboxylic acid synthesized in the skin, liver, and brain, and the first intermediate in L-histidine catabolism via histidase. It is a major UVB chromophore and natural moisturizing factor in the stratum corneum [88].<br>• Elevated levels associate with immunomodulation (e.g., NK cell inhibition), skin cancer risk, multiple sclerosis, and atopic dermatitis [89]. Low levels link to histidinemia (histidase deficiency) [90].<br>• Sweet spots were identified across all biological age acceleration quantiles for both sexes. This suggests that the metabolite level must be high enough to maintain epidermal barrier function and histidine metabolism, yet low enough to avoid the immunosuppressive and carcinogenic risks associated with excessive systemic accumulation. |
| S1E | 1,2-dilinoleoyl-GPC (18:2/18:2) (HMDB0008138) | Lipid | • A major membrane phospholipid species important for membrane fluidity and a reservoir for lipid mediators [38,91].<br>• Reduced circulating GPC(18:2/18:2) has been reported in squamous cell lung carcinoma versus controls, consistent with phosphatidylcholine remodeling and tumor-associated lipid metabolism [92]. Experimental work [91] also shows that exogenous 1,2-dilinoleoyl-GPC can increase insulin sensitivity in palmitate-treated myotubes and promote lipolysis in adipocytes.<br>• Sweet spots were found across all quantiles in both sexes. These optimal ranges likely represent the concentration required to support robust insulin signaling and membrane structural integrity, while avoiding the metabolic shifts characterized by aberrant phosphatidylcholine remodeling seen in chronic disease states. |
| S1F | 1-(1-enyl-palmitoyl)-2-palmitoleoyl-GPC (P-16:0/16:1)* (HMDB0011207) | Lipid | • A plasmalogen phosphatidylcholine with vinyl-ether linkage at sn-1 (plasmalogen 16:0) and ester-linked palmitoleic acid (16:1 n-7) at sn-2. It is enriched in heart, brain, and immune cells as a membrane structural lipid and endogenous antioxidant via its reactive oxygen species (ROS)-sensitive vinyl-ether bond [93].<br>• Elevated levels positively associate with increased percentage of fat in the liver and pancreas (proton density fat fraction [PDFF]) [94]. Low circulating plasmalogens including ethanolamine and choline species associate with type 2 diabetes (T2D), Alzheimer's disease, peroxisomal disorders, NASH [95].<br>• Sweet spots were found across all quantiles and sexes. In the context of oxidative stress, these sweet spots represent an optimal antioxidant buffer. Maintaining levels within this window may ensure sufficient protection against ROS without reaching the thresholds associated with ectopic fat deposition and metabolic syndrome. |
| S1G | 1-(1-enyl-stearoyl)-2-arachidonoyl-GPE (P-18:0/20:4)* (HMDB0005779) | Lipid | • A plasmalogen phosphatidylethanolamine with vinyl-ether 18:0 at sn-1 and arachidonoyl (20:4 n-6) at sn-2. Enriched in cardiovascular/nervous system membranes as antioxidant via ROS-labile vinyl-ether bond. Found in dairy, chicken and seafood [38,96].<br>• Higher levels show causal protective effect against peripheral arteriosclerosis and atherosclerotic cardiovascular disease (ASCVD) events [97]. Low plasmalogens associate with cardiometabolic diseases [98].<br>• Generally decreasing trends in all quantiles and sexes. Sweet spots were found for lower quantiles in Females, some quantiles in males. This suggests that for many, "more is better" due to its protective cardiovascular role. However, the identified sweet spots for certain groups suggest a saturation point where the marginal benefit for longevity plateaus or reverses. |

| Fig | Metabolite (HMDB) | Super pathway | Roles/related studies/identified associations |
|---|---|---|---|
| S1H | 1-oleoyl-2-arachidonoyl-GPE (18:1/20:4)* (HMDB0009069) | Lipid | • A diacyl glycerophosphoethanolamine with oleoyl (18:1 n-9) at sn-1 and arachidonoyl (20:4 n-6) at sn-2. It is a precursor for bioactive lipids including endocannabinoids via phospholipase actions; oleic acid from vegetable oils (olive/canola), arachidonic from animal fats/eggs [38].<br>• Elevated levels associate with increased coronary heart disease (CHD) risk [99]. low-moderate levels linked to reduced ASCVD risk via anti-inflammatory/endothelial effects.<br>• Sweet spots were found for lower quantiles for both sexes. The 0.9 quantiles in both sexes exhibited flatter trends compared to other quantiles. These results highlight the importance of maintaining the metabolite level within these ranges, likely through a balanced intake of plant and animal fats, specifically for those aiming to preserve their low biological age acceleration status. |
| S1I | 1-stearoyl-2-oleoyl-GPC (18:0/18:1) (HMDB0008038) | Lipid | • A diacyl phosphatidylcholine with saturated stearoyl (18:0) at sn-1 and monounsaturated oleoyl (18:1 n-9) at sn-2. It is abundant structural membrane phospholipid essential for lipid bilayer fluidity, lipoprotein assembly, and intracellular signaling precursor [93].<br>• Elevated circulating levels positively associate with liver fat content (PDFF) and hypercholesterolemia [94].<br>• Sweet spots were present across all biological age acceleration quantiles. This indicates that while this lipid is essential for membrane architecture, exceeding the identified optimal range likely marks a transition toward dyslipidemia and hepatic fat accumulation, thereby accelerating biological aging. |
| S1J | 4-hydroxy-2-oxoglutaric acid (HMDB0002070) | Lipid | • A key intermediate in the hydroxyproline degradation pathway. It is a substrate for mitochondrial 4-hydroxy-2-oxoglutarate aldolase (HOGA1) which cleaves it to pyruvate and glyoxylate [93].<br>• High level may indicate 4-hydroxy-2-oxoglutarate (HOG) aldolase (HOGA) deficiency, which causes higher endogenous oxalate synthesis leading to calcium oxalate kidney stone disease [99]. The consequence of an unusually low level is not yet understood.<br>• Sweet spots were found across all quantiles and sexes. This suggests that the identified range represents the optimal throughput of the hydroxyproline pathway, and levels outside this range likely signal metabolic bottlenecks that increase the risk of renal calcification and associated biological aging. |
| S1K | glycolithocholate sulfate* (HMDB0002639) | Lipid | • A sulfated glycine conjugate of lithocholic acid, which is a secondary bile acid produced by the bacterial action on chenodeoxycholic acid in the gut. Sulfation is a major detoxification pathway for bile acids in the liver, increasing their water solubility to facilitate excretion and prevent the accumulation of potentially toxic hydrophobic bile acids [38,100].<br>• Elevation associated with chronic hepatitis and cirrhosis [100,101].<br>• Despite generally decreasing trends, sweet spots were identified across all quantiles. This suggests that while efficient sulfation and excretion are generally signs of health, there is a specific range that reflects an optimal balance between microbial production and hepatic detoxification, minimizing biological aging. |

| Fig | Metabolite (HMDB) | Super pathway | Roles/related studies/identified associations |
|---|---|---|---|
| S1L | Glycoursodeoxycholate (HMDB0000708) | Lipid | • A glycine-conjugated secondary bile acid derived from ursodeoxycholic acid (UDCA) via hepatic conjugation and bacterial 7β-epimerization of chenodeoxycholic acid in the colon [93].<br>• Low serum/stool levels associate with type-2 diabetes, obesity, and metabolic dysfunction.<br>• While exhibiting generally decreasing trends, sweet spots were found across all biological age acceleration quantiles in both sexes. This result suggests that while metabolically protective, there is a specific range that most effectively slows biological aging. Falling below this range may signal dysbiosis or metabolic syndrome, while exceeding it could indicate an altered bile acid pool that no longer provides marginal longevity benefits. |
| S1M | Maleate (HMDB0000176) | Lipid | • A dicarboxylic acid structurally analogous to fumarate; it is not an endogenous human metabolite but a xenobiotic converted by microbial maleate hydratase to D-malate [93].<br>• Elevated levels (experimental exposure) associate with acute kidney injury, renal tubular cell dysfunction, and Fanconi syndrome [102].<br>• While no distinct sweet spot was identified, the PD plots revealed a "safe zone" with sharp non-linear increases in biological aging beyond a specific concentration. This underscores the microbiome's role in aging. While the body can tolerate a specific window of microbial maleate, exceeding this threshold appears to trigger rapid physiological decline, likely through renal stress. |
| S1N | oleoylcarnitine (C18:1) (HMDB0005065) | Lipid | • A long-chain acylcarnitine formed by esterification of oleic acid (18:1 n-9) with carnitine, facilitating mitochondrial transport of long-chain fatty acids for β-oxidation [93].<br>• Elevated plasma levels associate with diseases including schizophrenia, liver cirrhosis, cardiovascular mortality in dialysis patients, insulin resistance, and chronic heart failure [103]. A low level is associated with coronary heart disease [104].<br>• The presence of sweet spots across all quantiles reflects the metabolic necessity of efficient fatty acid oxidation. Concentrations below the sweet spot may indicate impaired energy production, while concentrations above it likely reflect mitochondrial overload or incomplete oxidation, resulting in biological age acceleration. |
| S1O | adipoylcarnitine (C6-DC) (HMDB0061677) | Lipid | • A medium-chain dicarboxylic acylcarnitine formed by esterification of adipic (hexanedioic) acid with L-carnitine. Like other acylcarnitines, its role is to shuttle dicarboxylic acyl groups into mitochondria for β-oxidation and energy production [93].<br>• Elevated adipoylcarnitine is observed in severe COVID-19 and other states of impaired oxidative metabolism [105].<br>• While there was no clear physiological sweet spot, the PD plot reveals a distinct "safe zone" characterized by a sharp, nonlinear increase in aging rates outside of a specific concentration window. |
| S1P | 1-stearoyl-2-linoleoyl-GPE (18:0/18:2) (HMDB0008994) | Lipid | • A diacyl glycerophosphoethanolamine featuring stearic acid (18:0) at sn-1 and linoleic acid (18:2n-6) at sn-2 positions. Contributes to cell membrane lipid bilayers, supporting fluidity, metabolism, and signaling as a major zwitterionic phospholipid.<br>• Downregulated in high/moderate vs. low cognitive impairment (ADAS-Cog) groups, suggesting links to Alzheimer's disease progression [106]. |

| Fig | Metabolite (HMDB) | Super pathway | Roles/related studies/identified associations |
|---|---|---|---|
| | | | • Sweet spots were found across all biological age acceleration quantiles in both sexes. |
| S1Q | pyridoxate (HMDB0000017) | Vitamins | • A methylpyridine derivative and primary urinary catabolite of vitamin B6 (pyridoxine/pyridoxal/pyridoxamine), formed via aldehyde oxidase and microbial pyridoxal 4-dehydrogenase. It reflects vitamin B6 status and excretion [93], It is one of the few water-soluble vitamins that can reach toxic levels, primarily because it can accumulate in the tissues and nerves over time. Paradoxically, the toxicity mimics Vitamin B6 deficiency [107,108].<br>• Low level of the metabolite is a direct signal of an underlying Vitamin B6 deficiency. Low Vitamin B6 has been strongly correlated with cognitive decline and an increased risk of developing cardiovascular disease [109] and neurodegenerative conditions like Alzheimer's disease [110].<br>• Sweet spots were found across all biological age acceleration quantiles in both sexes. Given its role as a catabolite, the sweet spot identifies the "Goldilocks" zone of Vitamin B6 status: high enough to support neuroprotection and cardiovascular health, but below the threshold where tissue accumulation triggers toxic effects on the nervous system. |
| S1R | erythritol (HMDB0002994) | Xeno-biotics | • A four-carbon sugar alcohol endogenously produced via pentose phosphate pathway (PPP) in erythrocytes, liver, kidney using ADH1/SORD enzymes from erythrose. Also dietary (wine, sake, beer, watermelon, pear, grape, soy sauce) and food additive absorbed by passive diffusion [93].<br>• Elevated plasma/urine levels associate with cardiovascular diseases [111] and type 2 diabetes risk [112].<br>• Sweet spots were found across all biological age acceleration quantiles in males. The health deterioration after this zone occurred quicker in the lower quantiles than higher quantiles. This highlights that for individuals with low baseline biological age, maintaining erythritol within a narrow range is critical, as they may be more sensitive to the metabolic disruptions associated with high erythritol levels. |
| S1S | stachydrine (HMDB0004827) | Xeno-biotics | • A proline derivative with N,N-dimethylpyrrolidinium structure, classified as a compatible osmolyte and secondary metabolite. Highest concentrations in capers (Capparis spinosa), detected in soybeans, limes. Potential biomarker for citrus/caper intake [93], and It has been studied for its potential health benefits [113,114].<br>• Elevated levels stachydrine exhibits cardioprotective effects via multiple pathways encompassing anti-inflammatory, antioxidant, anti-apoptotic, and modulation of calcium handling functions [115].<br>• PD plots reveal a general inverse correlation between stachydrine levels and biological age acceleration, supporting its reported health benefits. Notably, the identification of sweet spots suggests that there appears to be an optimal concentration range where stachydrine most effectively slows biological aging. This indicates that maintaining levels within this specific window, rather than simply increasing them indefinitely, may yield the greatest longevity benefits. |
